# CACNA1C as a Prognostic Biomarker and Therapeutic Target in High-Grade Serous Ovarian Cancer: Clinical Validation and Molecular Dynamics of Nifedipine Blockade

**DOI:** 10.64898/2026.04.19.719516

**Authors:** Mohamed A. Hammad, Kingsley Y. Wu, Eslam E. Abd El-Fattah, Karen S. Aboody, Chia-en A Chang

## Abstract

High-Grade Serous Ovarian Cancer (HGSOC) is the most lethal gynecological malignancy due to aggressive growth, widespread metastases, and high intra-tumoral heterogeneity. Poor prognosis is largely due to late diagnosis, hence there is an urgent need to identify novel biomarkers for screening, diagnosis, and monitoring. Here, we propose the voltage-dependent calcium channel hCaV1.2 encoded by CACNA1C as a potential biomarker and therapeutic target in HGSOC. Using IHC analysis for ten ovarian cancer patients, cytotoxicity assay, TCGA gene expression and survival analyses, homology modeling, molecular docking, Calcium channel membrane assembly and molecular dynamics simulations, we tested CACNA1C’s role in HGSOC progression and the effect of blocking on cancer cell survival.

We show that nifedipine (NIFE), a calcium channel blocker (CCB), had a tumor suppressive effect based on binding models predicted by three-dimensional computer assisted molecular modeling and in vitro validation using human HGSOC cell line. Using The Cancer Genome Atlas ovarian public cohort, we found CACNA1C mRNA expression strongly correlated with poor patient survival for late-stage and metastasis than primary. We also show strong correlation of CACNA1C protein expression using immunohistochemistry correlating with COH ovarian carcinomas patients’ disease progression. This research demonstrates that targeting HGSOC via CCBs may be therapeutically beneficial. By establishing further in vitro, in vivo, and clinical trials using FDA approved NIFE may be repurposed to target CACNA1C for HGSOC.

**Novelty and Impact:** High-grade serous ovarian cancer (HGSOC) remains lethal due to late diagnosis and drug resistance. This study identifies CACNA1C (Cav1.2) as a novel prognostic biomarker and therapeutic target in HGSOC, showing that elevated expression correlates with metastatic/recurrent disease and poor survival. Using molecular dynamics and in vitro models, we demonstrate that the FDA-approved calcium channel blocker nifedipine binds stably to Cav1.2 and suppresses tumor cell growth more effectively than cisplatin. These findings support repurposing nifedipine for biomarker-driven HGSOC therapy.

**Translational Relevance:** Late diagnosis and progressive relapses significantly contribute to the poor prognosis of ovarian cancer. Identification of a tumor biomarker that can be used for screening, diagnosis, and monitoring is critical for improving clinical outcome. Our findings demonstrate that CACNA1C is a viable diagnostic marker for HGSOC and that its blockade with CCBs reduces tumor progression, highlighting their therapeutic potential.

## 1. Introduction

Ovarian cancer (OC) ranks as the third most prevalent gynecologic malignancy globally, with 239,000 new cases each year, representing 152,000 deaths worldwide (1). It is the deadliest of all female reproductive cancers (2). OC risk increases with age, rising sharply after the age of 50, with the average age at diagnosis between 50 and 70 (3). Because the clinical manifestations of early OC are nonspecific or undetectable, OC is often called a silent killer, and approximately 70% of patients are diagnosed with an advanced stage (4). The 5-year survival rate for stage III is 34%, and only 15% for stage IV (5). However, current improvements in treatment methods have only slightly improved OC survival rates (6).

There are five subclasses of OC which include high-grade serous (70%), low-grade serous (< 5%), mucinous (3%), endometrioid (10%), and clear-cell (10%) carcinomas. OC is heterogeneous with different epidemiological and genetic risk factors, precursor lesions, patterns of spread, molecular events during oncogenesis, responses to chemotherapy, and prognosis (7). High-Grade Serous Ovarian Cancer (HGSOC) typically presents at advanced stage (III-IV) and, despite the initial response to surgical debulking and firstline therapy with carboplatin and paclitaxel (with or without bevacizumab), most tumors eventually develop drug resistance, with a 5-year survival generally below 30%. Therefore, there is an urgent need to clarify the potential molecular mechanisms of HGSOC and identify novel biomarkers for early diagnosis and prognosis assessment (8, 9).

There is abundant evidence demonstrating that calcium signaling plays an important role in cancer cell proliferation, apoptosis resistance, invasion, and drug resistance (10). Calcium channels are generally classified into two categories: voltage-gated calcium channels (VGCCs) and ligand-gated calcium channels (LGCCs) (11). VGCCs consist of multiple subunits, including subunit alpha1C (CACNA1C), coded by the α1 subunit Cav1.2 and is reported to be involved in regulating cell-matrix adhesion, collagen fibril organization, cell adhesion, cellular response to amino acid stimulus, and negative regulation of cell proliferation (12). Previous meta-analysis results showed that CACNA1C was upregulated in brain tumors, leukemia, breast cancer, and other tumors, suggesting its regulatory role in cancer progression (13). Bioinformatics analysis also revealed that mutations of CACNA1C was significantly associated with longer overall survival (OS) in endometrial cancer patients (14).

Topotecan, a topoisomerase I inhibitor, is commonly incorporated into chemotherapeutic regimens for the treatment of OC. Previous studies have reported that Topotecan can downregulate the expression of CACNA1C (15). Retinoids, derivatives of vitamin A, are well-documented for their anticancer properties, attributed to their high receptor-binding affinity and transcriptional regulatory functions (16). As adjuvant agents in OC therapy, retinoids have also been shown to reduce CACNA1C expression (17). Given the ad-verse effects associated with Topotecan and retinoids, the use of more selective calcium channel blockers is anticipated to offer improved therapeutic efficacy with reduced toxicity.

These data support the use of CACNA1C as a diagnostic biomarker for human cancers. The studies presented here aim to investigate the diagnostic value of CACNA1C in HGSOC and assess the effects of calcium channel blocking, using nifedipine (NIFE), on human A2780R, OVCAR8 and SKOV3 OC cell lines. The use of calcium channel blockers was associated with a reduced risk of developing serous ovarian carcinoma (18). We postulate its use as an adjuvant to traditional chemotherapeutic approaches, especially in the case of hypertension comorbidities.

## 2. Materials and Methods

### 2.1. OC patient FFPE sample for Immunohistochemistry

Ten ovarian cancer and matched normal ovarian tissue samples were acquired from consented patients under City of Hope (COH) IRB number 23599. Formalin-fixed paraffin-embedded tissue blocks were sectioned at a thickness of 4 μm and put on positively charged glass slides. IHC stains were performed on Ventana Discovery Ultra (Ventana Medical Systems, Roche Diagnostics, Indianapolis, USA) IHC Automated Stainer. Briefly, the slides were deparaffinized, rehydrated, and incubated with endogenous peroxidase activity inhibition reagents and antigen retrieval solution. The anti-human Cav1.2 polyclonal primary antibody (1:400; NOVUS BIOLOGICALS INC, CO, USA) was incubated and followed with DISCOVERY anti-rabbit HQ and DISCOVERY anti-HQ-HRP (Ventana) incubation. The stains were visualized with DISCOVERY ChromoMap DAB Kit and counterstained with hematoxylin (Ventana) and cover slipped. After IHC staining, whole slide images were acquired with a Ventana iScan HT Scanner (Roche) and viewed by iScan image viewer software.

### 2.2. Assessment of immunostaining

Images were scanned by a Hamamatsu nano zoomer scanner and analyzed using Visiopharm software. In each image we used custom APP to measure or count the area of an image analyzed, and the number of cells positive for each marker within that area. This quantitative information was further calculated.

### 2.3. Gene Expression and Survival Analyses using TCGA and public late-stage HGSOC patient databases

Whole-transcriptome sequences of 376 primary OC patients were downloaded from TCGA (TCGA-BRCA, https://portal.gdc.cancer.gov/projects/TCGA-BRCA). The “high” and “low” groups were segregated based on median mRNA expression values. Kaplan– Meier survival analysis was used to determine the survival differences between “high” and “low” mRNA expression groups, which were visualized by Kaplan–Meier plots and compared using Cox regression analysis, with P-values calculated by log-rank test using the Survival package in R. The survival differences were considered to be statistically significant when P-values were < 0.05.

Meta-analysis of ovarian transcriptome datasets in different cohorts of late-stage HGSOC patients (GSE98281, GSE101108, GSE102073, GSE133296, GSE141142, GSE149723, GSE155164) using QIAGEN OmicSoft Studio, OncoHuman_B38_GC33. The cohorts were further filtered by selecting ovarian late-stage patients, mRNAseq expression is higher in late stage metastatic and recurrent tumor (n=111) compared to primary tumor patients (n=263). Gene expression profiles of all tumor samples displayed as a dot plot. Boxes represent the median and the 25th and 75^th^ percentiles. Dots represent outliers.

### 2.4. Cell culture, chemicals and reagents

Nifedipine (NIFE) was obtained from Fujifilm Wako (Richmond, VA, USA). Cisplatin was obtained from Teva Pharmaceuticals (Parsippany, NJ). Dulbecco’s Modified Eagle Medium (DMED), Roswell Park Memorial Institute Medium (RPMI), and A2780R, OVCAR8 and SKOV3 were purchased from American Type Culture Collection (ATCC) (Manassas, VA, USA). Fetal bovine serum (FBS), and penicillin– streptomycin were purchased from Sigma (St. Louis, MO, USA). Iscove’s Modified Dulbecco’s Medium (IMDM) contains 4 mM L-glutamine, 4500 mg/L glucose, and 1500 mg/L sodium bicarbonate, and is supplemented with 10% (v/v) FBS, and 1% (W/V) antibiotics. Live A2780R, OVCAR8 and SKOV3 cells were incubated at 37°C in a 5% CO2 incubator (Thermoscientific, Waltham, MA, USA).

### 2.5. Cell treatment

The experiments were conducted with three replicates for each control or treatment group. A uniform cell density of 5 × 10^4^ cells/mL was maintained for all treatment and control cultures. Cells were treated with 312 µM of NIFE, CIS (19) or left untreated and incubated for 72 h at 37°C in a humidified 5% CO2 incubator (Thermoscientific, Waltham, MA, USA).

### 2.6. Cell proliferation assay

Cell viability was assessed with CellTiter 96° AQueous One Solution Cell Proliferation Assay kit from Promega (Madison, WI, USA) with few modifications. Briefly, 100 µL aliquots of treated or untreated cell suspensions, as described in the previous section, were seeded into 96-well polystyrene culture plates and 20 µL of assay reagent was added to each well. After 60 min of incubation at 37°C, the absorbance was read at 490 nm with a 96-well plate reader from BMG LABTECH GmbH (Ortenberg, Germany).

### 2.7. Homology Modeling of the Human Calcium Channel (hCaV1.2)

The 3D molecular structure of hCaV1.2 channel was built using the homology modeling method (20, 21). The recently reported cryo-EM structure of the α1 subunit of the rabbit CaV1.1 channel (PDB:6JP5) (22) was used as the template to build the hCav1.2 molecular structure in this study. In short, there are 37 sequence isoforms reported in the UniProt database (https://www.uniprot.org/uniprot/Q13936), and the isoform 1 (identifier: Q13936-1) was selected for the homology model construction. The BLAST search identified 50 models, and we selected the model rCaV1.1 having the sequence identity of 72.35% and with the highest Global Model Quality Estimation (GMQE) value (0.41) for the construction. The structure of the constructed hCaV1.2, comprising residues L105 to L1649 is depicted in SI Figure 1.

### 2.8. Atomistic Molecular Dynamics Simulations of the hCaV1.2-apo Structure

The constructed hCav1.2-apo molecular structure was subjected to energy minimization to clear possible clashes during the modeling step. Briefly, solvent molecules such as alcohol, DMSO *etc*., were removed during the system preparation. Missing hydrogens were added using Tleap implemented in the AMBER 20 package (23). The molecular structure was minimized in a three-step process: hydrogen, side chain, and the full protein, with 500, 1000 and 5000 steps, respectively. The resulting minimized structure was used in the following molecular docking calculation.

### 2.9. Molecular Docking Calculation of NIFE Compound in the hCaV1.2 Channel

The molecular docking calculation of the NIFE and the modeled hCaV1.2 structure was performed using Molecular Operating Environment (MOE) (24). The constructed hCaV1.2 molecular structure was selected from the energy-minimized step, as described in the previous section. The 3D dihydropyridine (DHP)-based structure, NIFE, was obtained from the PubChem database (ID:4485). During the molecular docking calculation, ligand binding sites were first searched by the implemented MOE-site finder tool using Alpha Shapes (25). There were 116 sites found from the searching step and three sites (#3, #21, and #73 among 116) within the pore of the hCaV1.2 were determined as the subsequent docking site (**SI Figure 2**). AM1BCC charge model (26) was used to assign charges for NIFE. The docking algorithm in MOE allowed induced fit and free-side chain rotation. London dG and GBVI/WSA dG scoring functions were selected. The resulting five binding modes from each site for the ligand with the lowest scorings were determined for the subsequent binding mode analysis (**SI Table 1**).

### 2.10. Calcium Ion channel membrane assembly

The CHARMM-GUI Membrane Builder was used to build the membrane system in both the apo-form and NIFE-bound structures prior to MD simulations (27-31). We selected binding mode 2 in site 21 as our structure from the docking result. The heterogeneous lipid bilayer was created by choosing palmitoyl-oleoylphosphatidylcholine (POPC) lipid with the ratio of x length and y length in 200 A□ in the inner and outer leaflets of the membrane. The systems were hydrated with 20.0 A□ thickness water on the top and lower leaflets. The system size along the X and Y dimensions are set as 190 A□ and 150 A□ for Z dimensions. We added an ionic concentration of 150 mM sodium chloride solution into the rectangle simulation box. We used Ligand Reader and Modeler to generate the parameter and force field for ligand NIFE (32). The fully assembled systems comprised about 513,000 atoms for both structures and were subject to NPT ensemble simulation setup at 300 K temperature. Parameters and inputs in the overall systems, including the calcium channel protein, lipid bilayer, water, ionic concentration, and ligand in the NIFE-bound structure were assigned using AMBER force field generator, and AMBER 14FFSB force field for protein and GAFF2 for ligand were selected in the force field options.

### 2.11. Classical MD Simulation

The MD simulation protocol generated by the CHARMM-GUI employs two consecutive constant temperature, constant volume (NVT) and constant temperature, constant pressure (NPT) equilibration steps after energy minimization. The minimization step included 2500 steps of steepest descent with 5000 minimization cycles, in which proteins, ligands, and lipid head groups were restrained. In the NVT equilibration step, simulations were performed with a time step at 1 fs for 250 ps, followed by NPT equilibration step with a time step at 2 fs for 2 ns. Heating of the systems from 50 K to 300 K with an increment of 50K was achieved in 200 ps (time step of 2 fs) at each temperature. Long-range electrostatic interactions were computed by the particle mesh Ewald method (33). The SHAKE constraint was applied all atoms including hydrogens (34). The Langevin thermostat with a damping constant of 2 ps^-1^ was applied to maintain a temperature of 300 K. A cut-off of 12 A was used for the vdW and short-range electrostatic interactions. All the simulations were performed under the periodic boundary conditions. The final production MD runs were performed under NPT ensemble for 200 ns (time step 2 fs), and each frame was collected at 10 ps time interval.

### 2.12. Statistical Analysis

All data were derived from ≥3 independent experiments. Statistical analyses were conducted using GraphPad Prism 9.4.1 (GraphPad Software, Inc., San Diego, CA). Values were calculated as mean ± standard error of the mean. Significant differences between the groups were determined using the unpaired oneway analysis of variance test. P<0.05 was considered to indicate a statistically significant difference.

## 3. Results

### 3.1 CACNA1C mRNA gene expression in HGSOC: TCGA data, patient cohort and IHC validation

To explore clinical associations of CACNA1C, a Meta-analysis of ovarian transcriptome datasets in different cohorts of late-stage HGSOC patients (GSE98281, GSE101108, GSE102073, GSE133296, GSE141142, GSE149723, GSE155164) was performed. Gene expression profiles of CACNA1C show higher expression in metastatic and recurrence patients compared to patients with primary cancer **(Fig, 1A)**, The high mRNA expression of CACNA1C is negatively correlated with overall survival with p-value= 0.1574; n=84 **(Fig.1B)**.

**1.**
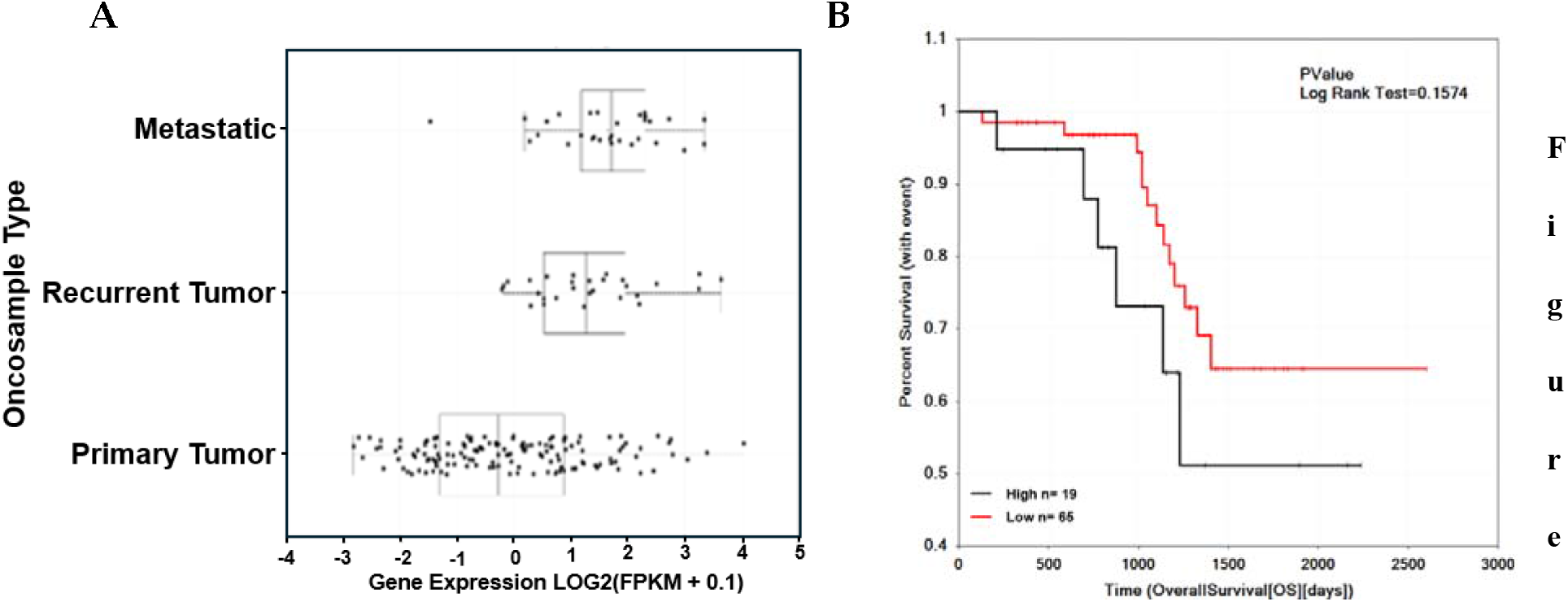
CACNA1C is a potential prognostic biomarker in late clinical stage using HGSOC late-stage ovarian cohort (n= 111). **(A)** patients’ mRNAseq expression shows higher expression in late clinical stage compared metastatic/recurrent to primary tumor using HGSOC patient cohort available cohort study. Clinical stage profiles of all tumor samples displayed as a dot plot. Boxes represent the median and the 25th and 75^th^ percentiles. Dots represent outliers. **(B)**. Kaplan-Meier survival analysis using HGSOC late-stage ovarian cohort shows as an unfavorable prognostic marker (*n*□= □84; p-value =0.158). CACNA1C mRNAseq expression values were converted into discrete variables by dividing the available population cohorts into “high CACNA1C” and “low CACNA1C” suggested by cutoff, *P*-value in the plot represents the result of log-rank test.

### 3.2. Clinical Characteristics and CACNA1C Expression in the COH Ovarian Cancer Cohort

Ten COH ovarian cancer patients were identified for genomic analysis of tumor samples, with key clinical characteristics shown in Table 1. 8 patients in this cohort were above 50 years of age with 6 identifying as white. Three patients presented with Stage I, two with stage II, two with stage III and one with stage IV HGSOC. The patients’ clinical stages range from 1c to 4b and included Asian, white, African American and one Hispanic or Latino patient, with an age range from 26 to 76 years old (**Table 1**). IHC for hCaV1.2 expression performed on OC tissue sections from a 10 patient COH cohort was 4.5 times greater than in paired normal ovarian tissues from the same patients (**Fig. 2**).

**Table 1:** HGSOC Patient characteristics.

| Patient serial # | Age at collection | Race | Cancer stage | % of IHC CACNA1C expression |
| --- | --- | --- | --- | --- |
| 01 | 70 | Asian | 1c | 12.6 |
| 02 | 61 | White | 3b | 8.2 |
| 03 | 56 | White | IC1 | 29.6 |
| 04 | 69 | Black or African American | NA | 35.4 |
| 05 | 50 | White | 2b | 9.0 |
| 06 | 50 | Asian | 4b | 15.6 |
| 07 | 76 | White | NA | 7.1 |
| 08 | 52 | White | 3b | 1.5 |
| 09 | 26 | Decline to Answer | 2 | 8.5 |
| 10 | 23 | White | 1c | 5.3 |

**Figure 2.**
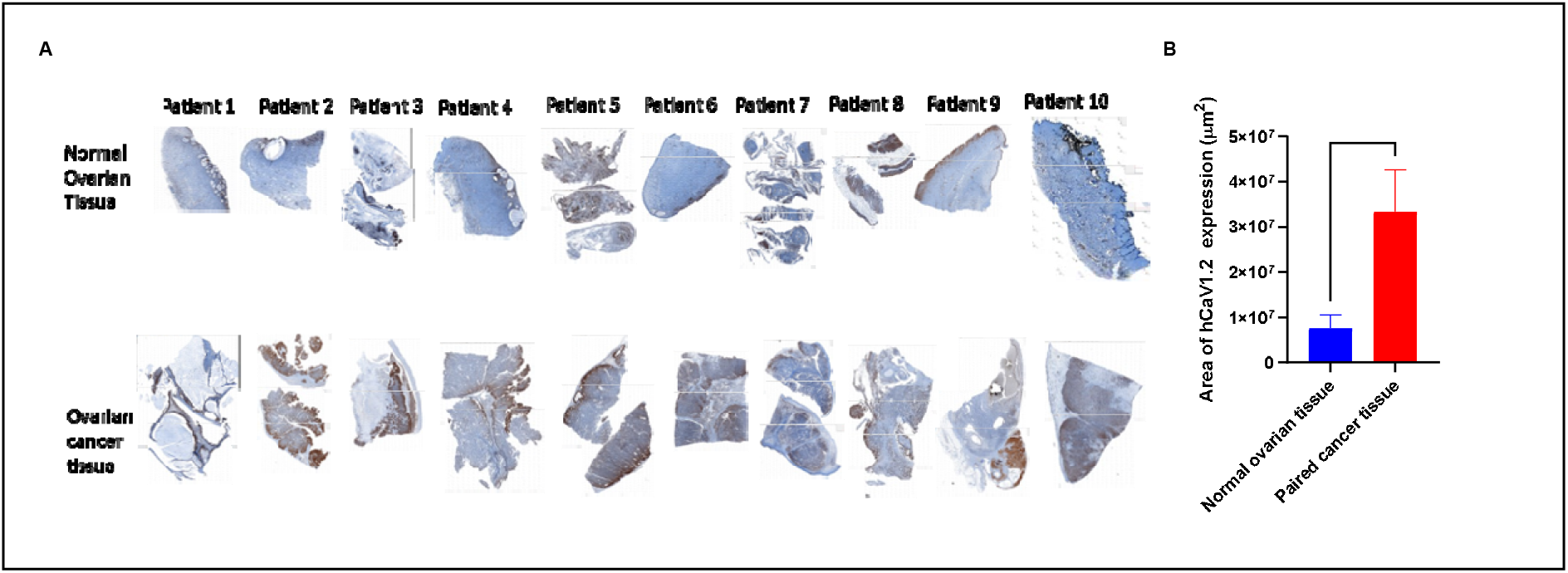
IHC of hCaV1.2 expression: **(A)** normal ovarian tissue (top row) paired OC tissue (bottom row), FFPE tissues were sectioned at a thickness of 4 μm, stained with anti-human Cav1.2 polyclonal primary antibody then IHC stains were performed on Ventana Discovery Ultra IHC Automated Stainer. (brown). **(B) Automated IHC measurements (hCaV1.2 %Pos area/slide)**

### 3.3. NIFE kills HGSOC Cells

Our cytotoxicity experiments showed that NIFE induced inhibitory effects on OC cell proliferation (**Fig. 3**). Notably, NIFE showed more cytotoxic effect than the standard of care Cisplatin (CIS) at the same dose (312 uM) on 3 different human OC cell lines (A2780R, OVCAR8 and SKOV3).

**Figure 3.**
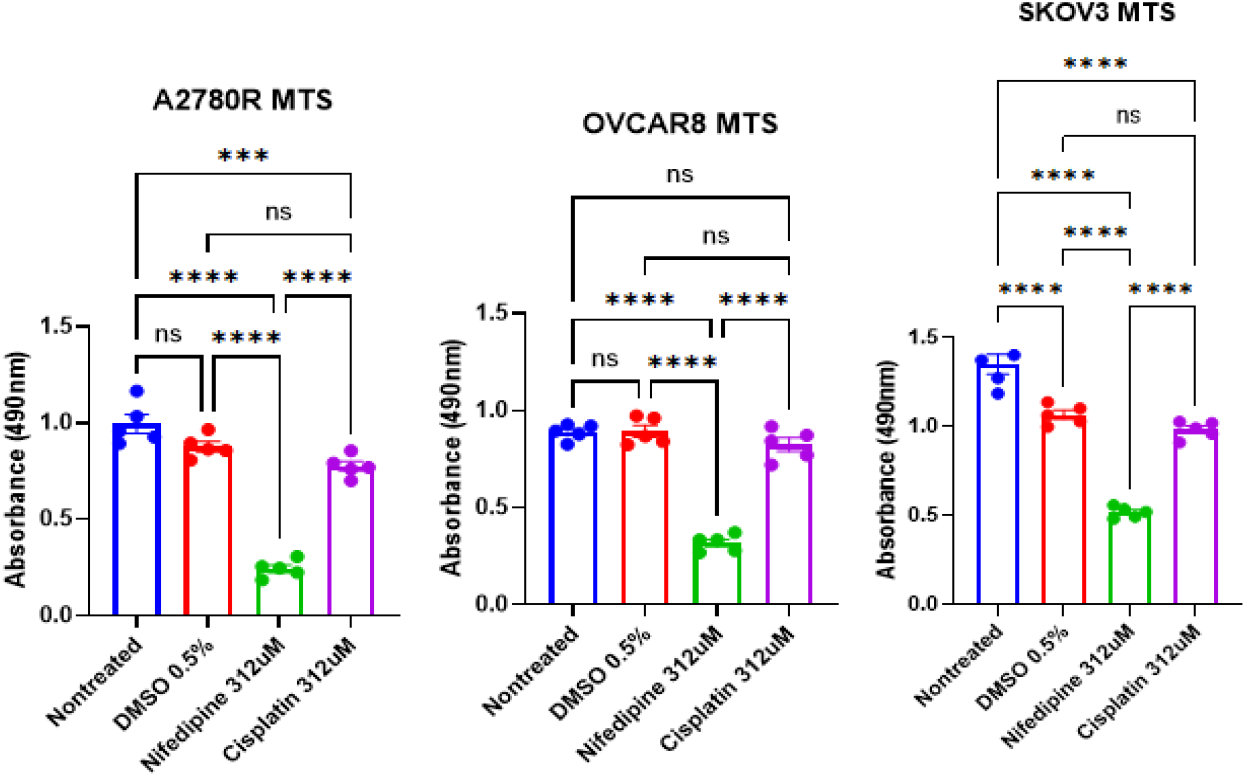
Cytotoxic effect of NIFE on human ovarian cancer cell lines (A2780R, OVCAR8 and SKOV3). Cells were treated with 312 µM of NIFE for 72h. Cell viability was tested by cell proliferation assay using a spectrophotometer at 490 nm. Results were expressed as means of three independent experiments ± standard error mean. *p-values < 0.05, **p-values < 0.01, ***p-values < 0.001, ****p-values < 0.0001 were considered statistically significant.

### 3.4. Molecular Docking Calculations of NIFE in hCaV1.2 Constructed Structure

To investigate and understand the binding mechanism of compounds in the human CaV1.2 channel, we applied molecular docking in the current study. Due to the lack of molecular structure information on the human CaV1.2 channel in the protein data bank, we built our hCaV1.2 molecular structure using homology modeling (20). The structure of the constructed hCaV1.2, comprising residues L105 to L1649 is depicted in **Figure 4**. The 3D dihydropyridine-based structure, NIFE, was obtained from the PubChem database and assigned partial charges using Austin Model 1-Bond Charge Correction (AM1BCC). To dock the ligand into the constructed hCaV1.2 model, we performed MOE-site finder to search for possible binding sites. Among a total of 116 searched sites, three sites (site #3, #21 and #73) located in the pore domain of the channel were considered for the subsequent docking study. Site #3 and #21 located in the fenestration enclosed by domains III and IV, except that site #3 covered a wider range in the pore domain. Site #73 located in the center cavity, slightly off the pore axis toward the fenestration formed by the domains III and VI. Our site-finder search method identified these three sites, which were also observed in other calcium channel studies (22, 35-38). Ligand docking calculations were performed in these three sites and the resulting five binding modes from each site for NIFE with the lowest scorings were determined for the subsequent binding mode analysis **(SI Table 1)**. Scores calculated from site #21 were the lowest among the three sites for the ligand.

**Figure 4.**
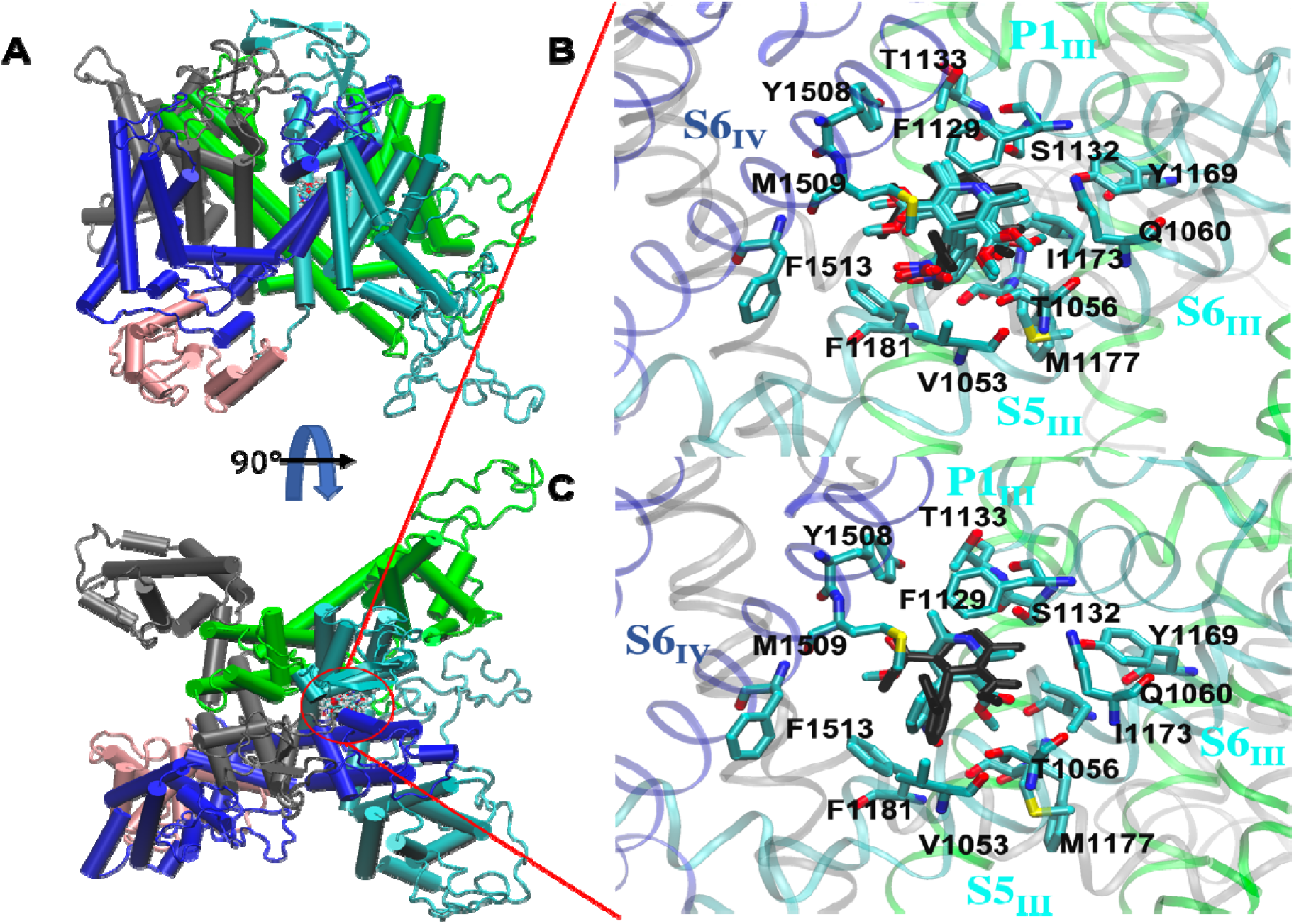
NIFE binds to the fenestration enclosed by domain III (cyan) and IV (blue) of hCaV1.2. **(A)** 15 binding modes were overlaid in the fenestration enclosed by domain III (cyan) and IV (blue). **(B)** Four binding modes were superimposed with the crystal structure (black) (PDB ID: 6JP5). **C**. One binding mode was superimposed with the crystal structure (black), nitrophenyl ring rotating 180°.

The overall binding modes based on the scoring consideration were subjected to further analysis **(Fig. 4)**. In the NIFE binding modes, all 15 poses (five from each site) are located in the fenestration enclosed by domains III and IV **(Fig. 4A)**. According to the previous reports and other crystal structures (22, 35), poses locating in the fenestration enclosed by domains III and IV, close to helical segments S5_III_, S6_III_, S6_IV_ and P1_III_ were further analyzed.

In the NIFE binding modes, four molecular poses (site #21 mode 2 and 3, site #73 mode 1 and 3, bold in Table S1) were found to closely agree with the reported crystal structure (PDB ID: 6JP5) **(Figure 4B)** (22). In the structure, the whole molecule was embedded in a hydrophobic core formed by Phe1129 and Thr1133 on P1_III_, Val1053 on S5_III_, Tyr1169, Ile1173, Met1177 and Phe1181 on S6_III_, and Tyr1508, Met1509 and Phe1513 on S6_IV_. The N1 amine in the dihydropyridine group formed a H-bond with the hydroxyl group of Ser1132 on P1_III_. Thr1056 and Gln1060 on S5_III_ each formed a H-bond with two oxygens from the C3-ester in the NIFE. In addition to the polar interactions, non-polar interactions were also observed; Phe1129 was in parallel with the dihydropyridine group, forming π-π interaction. The phenyl ring in the nitrophenyl group was T-shaped stacking to the Phe1181. Interestingly, we observed one binding mode (site #21 mode 1) **(Fig. 4C)** that was also overlaid with the crystal structure, except the nitrophenyl ring rotating 180°. The whole ligand molecule was still accommodated in the hydrophobic pocket as described above.

### 3.5. Atomistic Molecular Dynamics (MD) Simulations of the Apoprotein and NIFE-Bound Complex

To gain a better understanding of the NIFE interactions in the hCaV1.2 complex for further drug development, we performed MD simulations to examine protein conformational changes before and after NIFE binds. We, therefore, applied MD simulations for both NIFE-hCaV1.2 complex and apoprotein (hCaV1.2 protein only). Since the hCaV1.2 ion channel is a membrane-bound protein, the structure was embedded in a POPC bilayer with water molecules and ions (**Figure 5**) (See methods for details).

**Figure 5.**
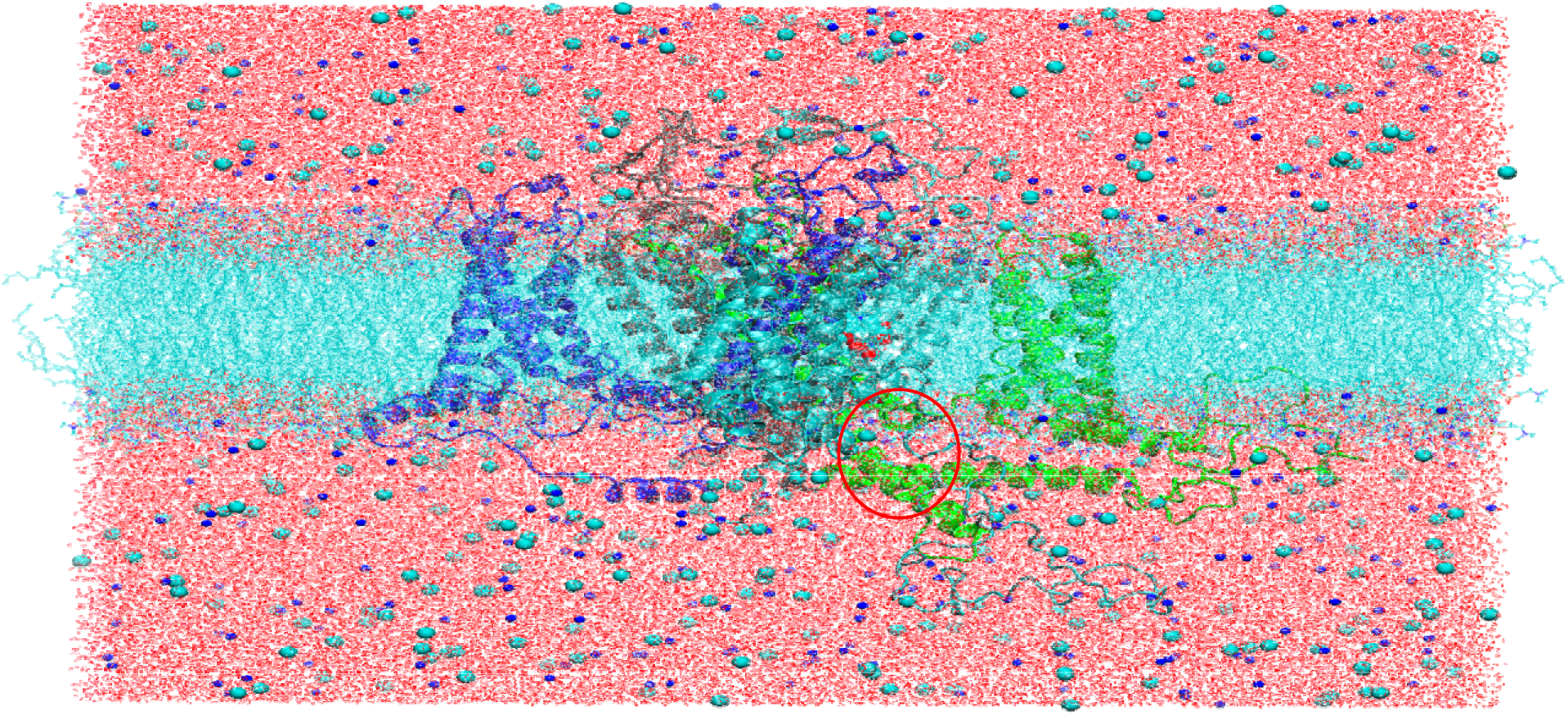
Snapshot of the constructed hCaV1.2 ion channel embedded in a POPC bilayer with water molecules, sodium, and chloride ions. Four subdomains (gray for domain I, green for domain II, cyan for domain III, and blue for domain IV) are shown. Ligand NIFE (VDW representation) is encapsulated between domains III and IV, which is circled in red in the figure.

The time-dependent root-mean-square-deviation (RMSD) results showed that apoprotein reached the equilibrium at 80 ns, whereas NIFE bound complex reached the equilibrium at 20 ns in the 200 ns MD simulations (**SI Fig, 3**). Inlet in **SI Figure 3** is the RMSD results for residues in the ligand bound pocket in both systems. It showed that NIFE bound pocket reached the equilibrium at 50 ns in the NIFE bound structure, but it took about 100 ns for the apoprotein to reach equilibrium. Our RMSD results implied that the NIFE bound structure stabilized and reached a local energy minimum quicker than the apoprotein structure, but two structures reached stability in our 200 ns MD simulations.

The exploration of the rotameric states of the residues provides useful information regarding protein flexibility and protein conformational changes and dynamics. To better understand the interactions between the NIFE and the residues, we applied a T-analyst program to study the residues’ rotameric states in the binding pocket in both the NIFE-hCaV1.2 complex and the apoprotein (39). We focused on the residues surrounding the NIFE binding pocket; Val1053, Phe1129, Ser1132, Phe1176, Met1178, Phe1181, Met1509, as these key residues were reported to be involved in mutagenesis experiments with known LTCC antagonists (38) (**Fig. 6**). The rotameric state of two key residues Phe1129 and Ser1132 on P1_III_ were reported and compared (**Fig. 6C, D** for apoprotein and **Fig. 6F,G** for NIFE bound structure). Phe1129 maintained one rotameric state in the whole MD simulations by forming a strong - interaction with the dihydropyridine ring in the NIFE, (Right panel, **Figure 6B and 6F**) while it was observed to flip its phenyl ring 90º in the apoprotein structure during the simulation. (Left panel, **Figure 6A** and **6C**).

**Figure 6.**
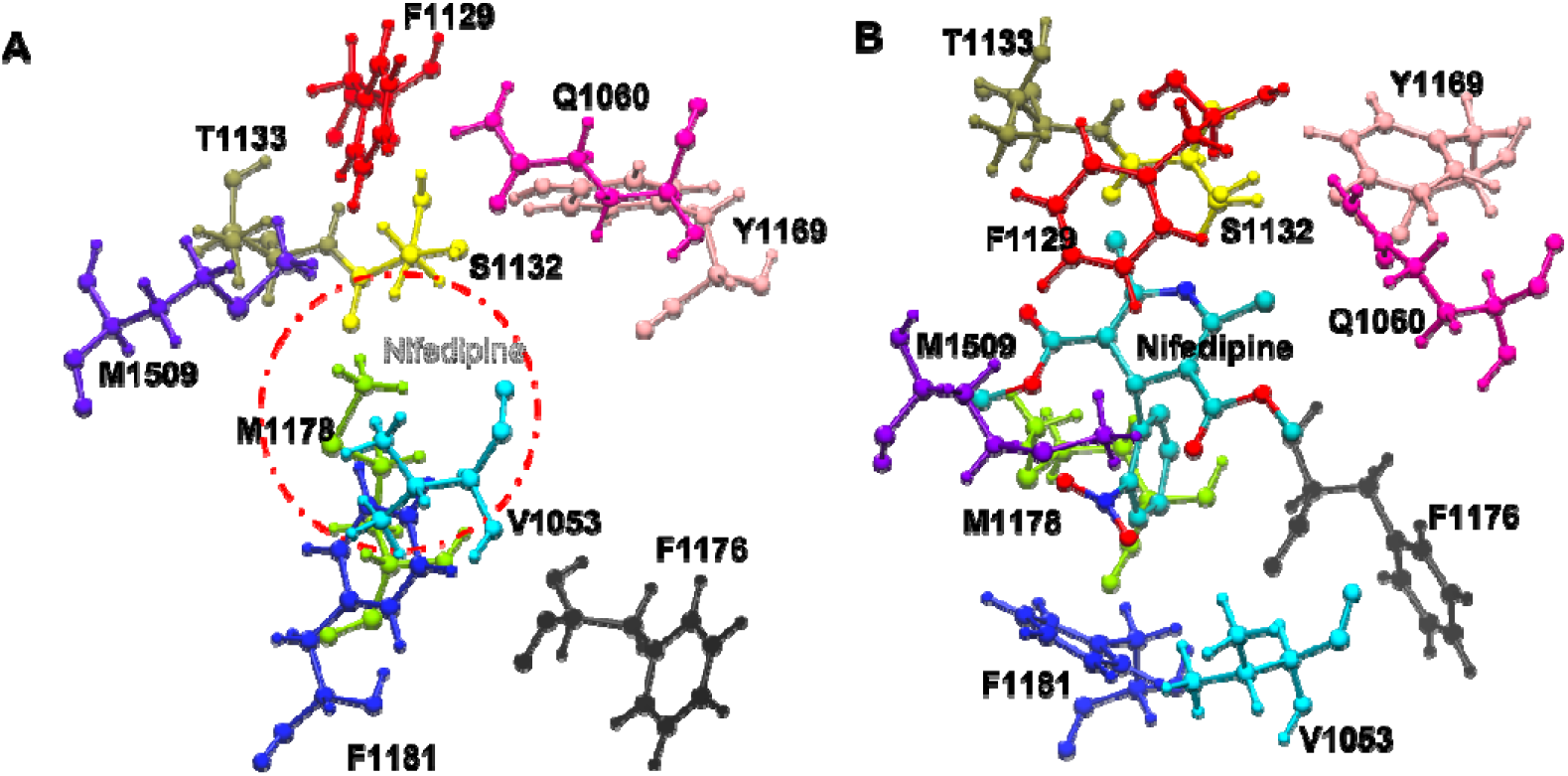
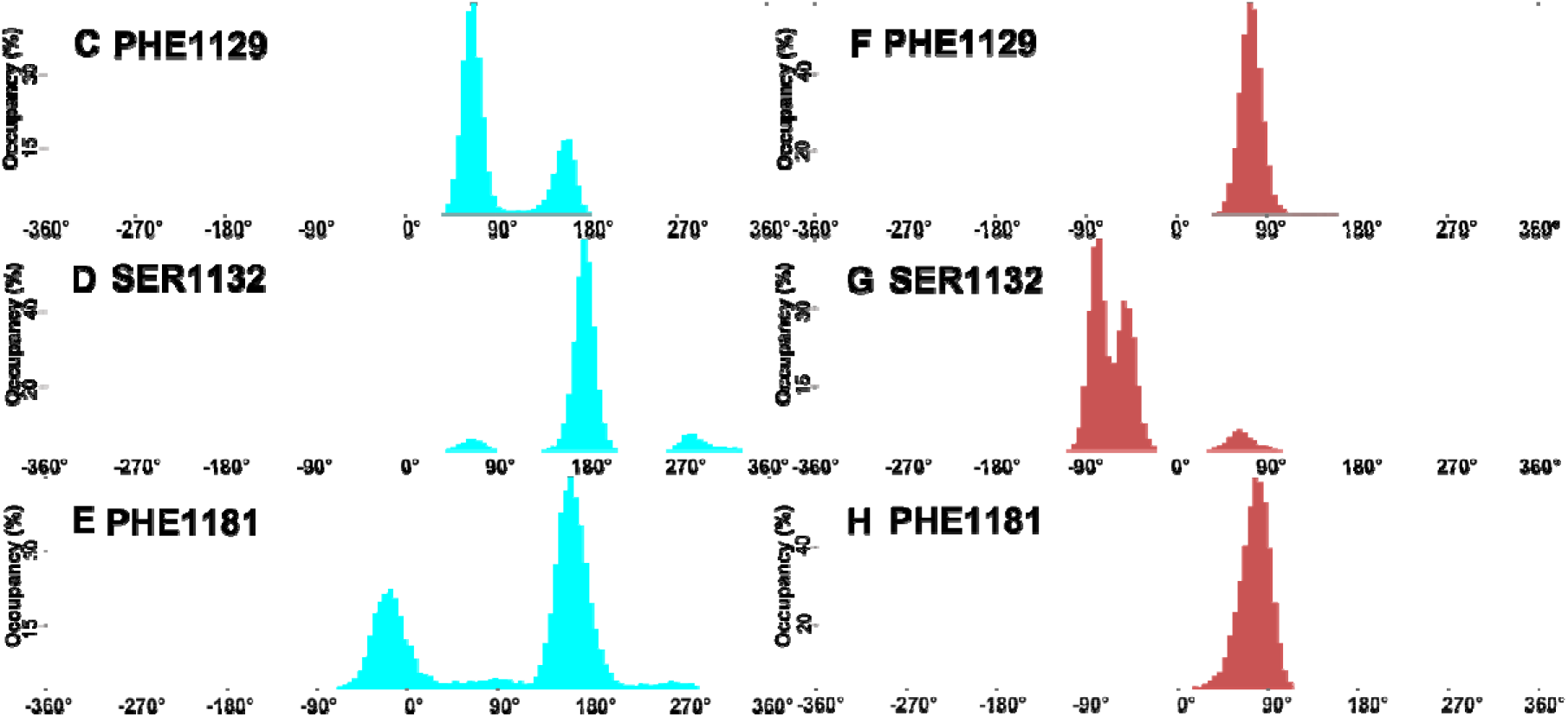
Snapshot of residues around the NIFE binding pocket in apoprotein structure **(A)** and NIFE bound structure **(B)**. The dash line circle in (A) indicates the position of the NIFE bound in the pocket. **(C)-(H)** Clustering conformations of residues surrounding the ligand binding pocket. Each histogram can represent one rotameric state. (C)/(F) (Phe1129), (D)/(G) (Ser1132), E/H (Phe1181). Left panel is for apoprotein structure and right panel is for NIFE bound structure.

Previous observations showed that Ser1132 has a strong binding affinity with the DHP antagonists (40, 41) hence, its hydroxyl group formed a stable H-bond with the N1 amine in the dihydropyridine group of NIFE in the NIFE bound structure, (**Fig. 6G**) but rotated away in the apoprotein structure, alternatively forming H-bonds with Glu1060 on S5_III_ and Try1169 on S5_IV_. (**Fig. 6D**) Phe1181 on S6_III_ formed a stable T-shaped aryl-aryl interaction with the phenyl ring in the nitrophenyl group of NIFE, as indicated by one rotameric state, (**Fig. 6H**) but rotated alternatively to fill the ligand binding pocket when NIFE was absent (**Fig. 6E**).

It was observed that there are three rotameric states of Val1053 in the apoprotein structure (**SI Fig. 4A**), whereas there was one rotameric state of Val1053 (**SI Fig. 4E**) in the NIFE bound structure, indicating that Val1053 was more flexible in the apoprotein as it lost its ligand bound coordination when ligand was absent. The hydrophobic residues Phe1176 and Met1178 on S6_III_ each had one rotameric state in the NIFE bound structure (**SI Fig. 4F and 4G**, respectively), as they were observed to be pushed outwards of the binding pocket with the presence of the NIFE. Two rotameric states were observed in these two residues when there was a vacancy in binding pocket in the apo structure (**SI Fig. 4B, C**).

Met1509 on S6_IV_ was also observed to rotate alternatively to fill the binding pocket in the apoprotein structure, (**SI Fig. 4D**) but was pushed outwards when NIFE was present. (**SI Fig. 4H**) In the NIFE bound state, these residues coordinated with the NIFE and stabilized the local environment in the binding pocket, while in the apoprotein structure, these residues are free to move and rotate since more free space is allowed in the binding pocket. Our investigation of hCaV1.2 ion channel dynamics suggested that NIFE interacted with the side chain of residues Val1053, Phe1129, Phe1176, Met1178, Phe1181 and Met1509 in the pocket via mainly the hydrophobic interactions.

In particular, the NIFE showed a strong H-bond interaction with Ser1132, alongside persistent *π-π* stacking interactions with the Phe1129 and Phe1181. These residues were able to coordinate with the NIFE, yet it was observed that it is highly flexible in the apoprotein structure, which may construct a pocket that binds to compounds sharing a similar scaffold as NIFE. This analysis provides useful information for further modifications of the NIFE and the discovery of new inhibitors targeting hCaV1.2 ion channel.

To explore the activation/inactivation state caused by the effect of the NIFE bound to the structure, we measured the permeation path of the hCaV1.2 ion channel in both the apoprotein and NIFE bound structures (**Fig. 7**). Previous studies have shown that DHP antagonists selectively inhibit L-type calcium channel under persistent depolarized conditions, which linked to sealed intracellular gate (42). We applied HOLE program (43) to calculate the radius of the calcium permeation path along the channel pore. Green dots indicated a widen intracellular gate and selective filter in the apoprotein, whereas red dots indicated a narrow intracellular gate and selective filter. It is suggestive that the channel pore is less permeable for calcium ions to pass through since the NIFE bound structure exhibits depolarized conformation and is in inactivated state.

**Figure 7.**
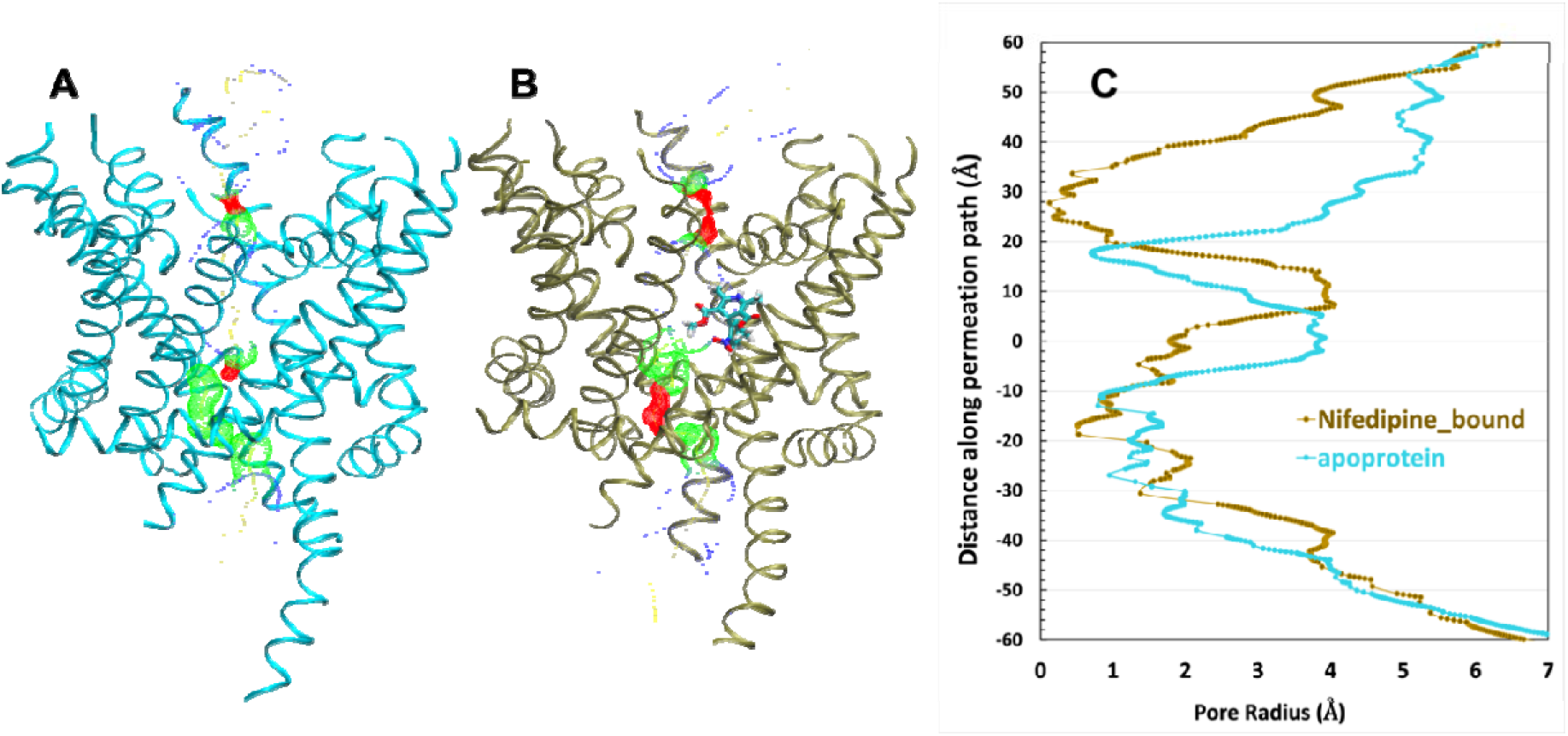
The permeation path of hCaV1.2 apoprotein (A) and NIFE bound (B) structures calculated by HOLE(43) are illustrated by dots in the panel. The pore radii <1.15 Å are colored in red, between 1.15 Å and 2.3 Å are colored in green and > 2.3 Å are colored in blue. Measurement is according to (22) (C) Pore radii along the permeation path.

## 4. Discussion

HGSOC is the most prevalent and lethal subtype of epithelial ovarian cancer, with limited therapeutic options and poor prognosis. Due to late-stage diagnosis attributable to nonspecific or absent symptoms, HGSOC has one of the worst outcomes among gynecological cancers. Currently, HGSOC detection relies on conventional techniques such as transvaginal ultrasound and carbohydrate antigen 125 (CA125) testing; however, combined use of these modalities has not resulted in appreciable reductions in overall mortality rates.

The current paradigm for managing HGSOC has therefore shifted toward early diagnosis through liquid biopsies, a non-invasive method of detecting molecular markers, underscoring the urgent need for improved biomarkers to enable more rapid and accurate diagnosis, monitor disease progression during chemotherapy, and predict recurrences.

In the present study, our TCGA analysis identified CACNA1C, which encodes the L-type voltage-gated calcium channel Cav1.2, as differentially expressed between normal ovarian and ovarian cancer transcriptomes. Importantly, our analysis of ovarian tumor tissue from patients with ovarian cancer revealed that Cav1.2 protein expression was significantly increased compared to matched normal ovarian tissue from the same patients, and this elevated expression was negatively correlated with survival rate. These findings are consistent with broader evidence from TCGA ovarian cancer cohorts demonstrating that high CACNA1C expression independently predicts poor overall survival (HR ∼1.58), shorter platinum-free intervals, and enrichment in platinum-resistant recurrent disease (44).

The mechanistic basis underlying the association between CACNA1C overexpression and poor clinical outcomes in HGSOC likely involves multiple oncogenic pathways. Previous studies have demonstrated that elevated Cav1.2 activity promotes sustained intracellular calcium signaling that activates NFAT, NF-𝒦B, and CaMKII pathways, leading to enhanced proliferation, survival, and chemotherapy resistance (45-47).

Furthermore, changes in CACNA1C expression have been shown to correlate with promotion of epithelialmesenchymal transition (EMT), increased invasive capacity, and enrichment of cancer stem cell populations—mechanisms that might facilitate the peritoneal dissemination characteristic of HGSOC (48).

While Li et al. (48) reported a favorable prognostic association for CACNA1C in ovarian cancer, their analysis encompassed heterogeneous histological subtypes. In contrast, our study focused exclusively on highgrade serous ovarian carcinoma (HGSOC), which represents a molecularly and clinically distinct entity with unique genomic alterations, including near-universal TP53 mutations and distinct origins from fallopian tube epithelium. Our meta-analysis of seven independent HGSOC-specific transcriptomic datasets (GSE98281, GSE101108, GSE102073, GSE133296, GSE141142, GSE149723, GSE155164) consistently demonstrated elevated CACNA1C expression in metastatic and recurrent disease compared to primary tumors, suggesting subtype-specific prognostic implications and a potential oncogenic role in disease progression rather than tumor initiation, an association that may not have been captured in studies analyzing heterogeneous ovarian cancer cohorts at variable disease stages.

These findings were substantiated at the protein level through immunohistochemical analysis, which demonstrated a 4.5-fold increase in hCaV1.2 expression in HGSOC tissues compared to patient-matched normal ovarian tissues across a diverse 10-patient cohort spanning multiple ethnicities and disease stages. The therapeutic relevance of CACNA1C overexpression in HGSOC was further supported by our cytotoxicity studies demonstrating that the L-type calcium channel blocker nifedipine exhibited superior cytotoxic effects compared to the standard-of-care agent cisplatin at equivalent doses across three HGSOC cell lines (A2780R, OVCAR8, and SKOV3), while our molecular docking and atomistic MD simulations revealed stable binding of nifedipine within the hCaV1.2 channel pore, promoting a depolarized, inactivated channel state that provides a structural rationale for targeting CACNA1C-overexpressing HGSOC tumors with dihydropyridine-based therapeutics. Although the negative correlation between CACNA1C expression and overall survival in our HGSOC cohort did not reach statistical significance (p=0.1574, n=84), the consistent trend across multiple independent datasets, combined with protein-level validation and functional therapeutic data, suggests a biologically meaningful association, and larger prospectively designed studies focusing specifically on HGSOC are warranted to definitively establish the prognostic significance of CACNA1C in this aggressive subtype.

Additionally, CACNA1C overexpression has been implicated in driving multi-drug resistance phenotypes through enhanced drug efflux via MDR1/P-glycoprotein upregulation, impaired drug-induced apoptosis, and activation of DNA repair machinery, conferring resistance to platinum agents, taxanes, and potentially PARP inhibitors. Our findings demonstrating the negative correlation between Cav1.2 expression and patient survival are therefore consistent with these established mechanisms and support the role of CACNA1C as a driver of aggressive tumor behavior.

To functionally validate the correlation between Cav1.2 activity and tumor progression, we tested the effect of the calcium channel blocker nifedipine on ovarian cancer progression. Notably, our results demonstrated that blocking Cav1.2 channels inhibited cancer growth significantly more than the traditional chemotherapeutic agent cisplatin, suggesting that calcium channel blockade may represent a viable therapeutic strategy for HGSOC. To further confirm that the inhibitory effect of nifedipine on cancer growth is primarily mediated through calcium channel blockade, we performed homology modeling of the human calcium channel (hCav1.2), molecular docking simulations, and calcium ion channel membrane assembly analysis. Our computational analyses revealed that nifedipine interacted with residues in the binding pocket primarily via hydrophobic interactions, providing structural evidence supporting the functional role of calcium channels in ovarian cancer growth and validating our experimental findings.

## 5. Conclusions

In conclusion, our research demonstrates that CACNA1C can serve as a novel biomarker and potential therapeutic target for individuals with HGSOC, functioning both as a mechanistic driver of tumor aggressiveness and an independent prognostic indicator. The significant differential expression of CACNA1C between normal ovarian and ovarian cancer transcriptomes, coupled with the negative correlation between Cav1.2 protein expression and patient survival, supports its utility for HGSOC diagnosis, progression monitoring, and prediction of clinical outcomes. Furthermore, our findings suggest that nifedipine, a calcium channel blocker with an established safety profile, emerges as a promising FDA-approved drug that can be repurposed as a therapeutic option for HGSOC, given its demonstrated ability to inhibit cancer growth more effectively than conventional chemotherapeutic agents such as cisplatin. However, further validation using larger HGSOC cohorts from City of Hope and publicly available databases is warranted, combining tissue imaging with multi-omic integration analysis to comprehensively investigate the molecular pathways connected to CACNA1C in ovarian cancer. Such efforts will create a productive framework for developing more potent and tumor-selective calcium channel blockers while establishing the foundation for biomarker-driven clinical trials evaluating CACNA1C antagonism in combination with standard chemotherapy, ultimately offering new hope for patients with this aggressive malignancy.

DSC traces of (A) GUV suspension (55 µM total lipid; 43 µg/mL) prepared with doubled lipid deposition relative to microscopy samples to improve signal-to-noise and (B) MLV suspension (1 mg/mL); both samples were prepared in 100 mM sucrose. Heating rate: 1 K/min. The main transition temperature is ∼31.9 °C, approximately 9 °C below the T_m_ of pure DPPC (Fig. S8). Note that the GUV profile may contain contributions from non-unilamellar material and is expected to represent a lower bound on the true breadth of the gel-fluid transition in isolated GUV bilayers (see main text). Colored curves represent five consecutive heating scans, with the averaged profile shown as a thick grey line.

## Abbreviations

CCB: calcium channel blocker
CIS: cisplatin
COH: City of Hope
DHP: dihydropyridine
EMT: epithelial–mesenchymal transition
FFPE: formalin-fixed paraffin-embedded
HGSOC: high-grade serous ovarian cancer
IHC: immunohistochemistry
LGCC: ligand-gated calcium channel
MD: molecular dynamics
MOE: Molecular Operating Environment
NIFE: nifedipine
OC: ovarian cancer
OS: overall survival
PDB: Protein Data Bank
POPC: palmitoyl-oleoyl-phosphatidylcholine
RMSD: root-meansquare-deviation
TCGA: The Cancer Genome Atlas
VGCC: voltage-gated calcium channel

## Author Contributions

Conceptualization, K.Y.W., C.A.C. and M.A.H.; methodology, K.Y.W., C.A.C. and M.A.H.; validation, C.A.C. and M.A.H.; formal analysis, K.Y.W., C.A.C. and M.A.H.; investigation, K.Y.W., C.A.C. and M.A.H.; data curation, K.Y.W., C.A.C. and M.A.H.; writing—original draft preparation, K.Y.W., C.A.C., EEAE and M.A.H.; writing, editing— K.Y.W., C.A.C, K.S.A., E.E.A. and M.A.H.; supervision, C.A.C. and M.A.H. ; project administration, K.S.A., K.Y.W., C.A.C. and M.A.H.; funding acquisition, C.A.C. All authors have read and agreed to the published version of the manuscript.

## Data Availability Statement

The data presented in this study are available on request from the corresponding author. Conflicts of Interest: All authors declare no conflict of interest. The funders had no role in the design of the study; in the collection, analyses, or interpretation of data; in the writing of the manuscript, or in the decision to publish the results.

## Acknowledgements

Research reported in this publication included work performed in the Pathology Core supported by the National Cancer Institute of the National Institutes of Health (NIH) under grant number P30CA033572. The computation work was supported by the NIH (R01GM109045 to CAC). The content is solely the responsibility of the authors and does not necessarily represent the official views of the National Institutes. The authors would like to acknowledge the COH Center for Informatics most notably Translational Bioinformatics, and the utilization of the POSEIDON platforms for data exploration, visualization, analysis, support, and training. We thank Dr. Alemayehu Gorfe for insightful discussion and suggestions.

## Disclaimer

The content is solely the responsibility of the authors and does not necessarily represent the official views of the National Institutes of Health or any other funding agency.

## Ethics Statement

Ovarian cancer ad matched normal ovarian tissue samples were acquired from consented patients under City of Hope (COH) IRB number 23599. All procedures followed the ethical standards of the institutional research committee and the Helsinki Declaration.

## Supplemental Information (SI): Figures and Table

**SI Figure 1.**
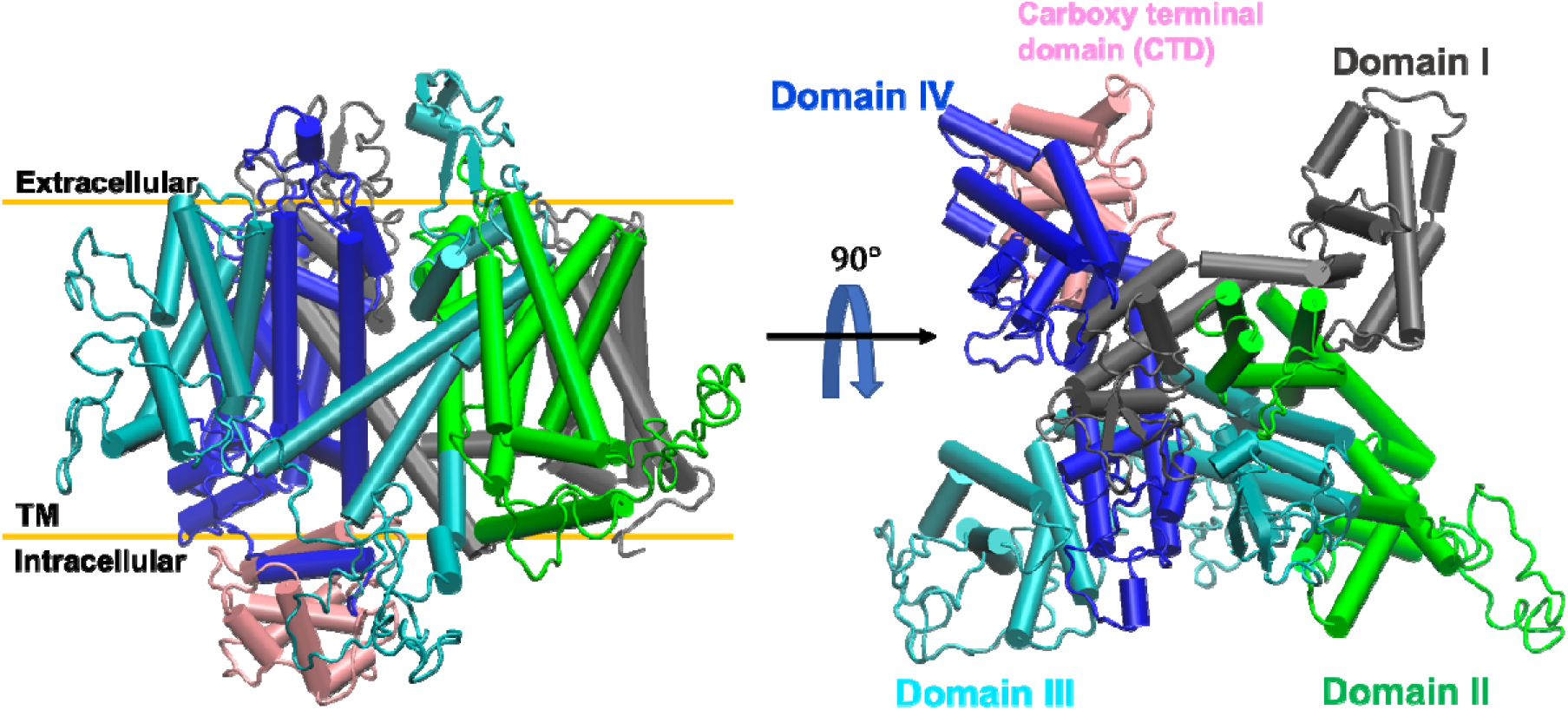
Structure of the constructed hCaV1.2 channel. Structure of the hCaV1.2 channel generated by homology modeling from lateral view (left) and rotate 90° from extracellular view (right). Scheme of cartoon representation for hCaV1.2 structure with four domains and carboxy terminal domains are indicated; domain I in gray, domain II in green, domain III in cyan, and domain IV in blue. Each domain is composed of six transmembrane helices (S1-S6), where the first four, S1-S4, formed the voltagesensing domain (VSD) while the other two, S5-S6, formed the pore domain (PD). The extracellular segment that connects helices S5 and S6 formed two re-entrant short helices denominated P1 and P2.

**SI Figure 2.**
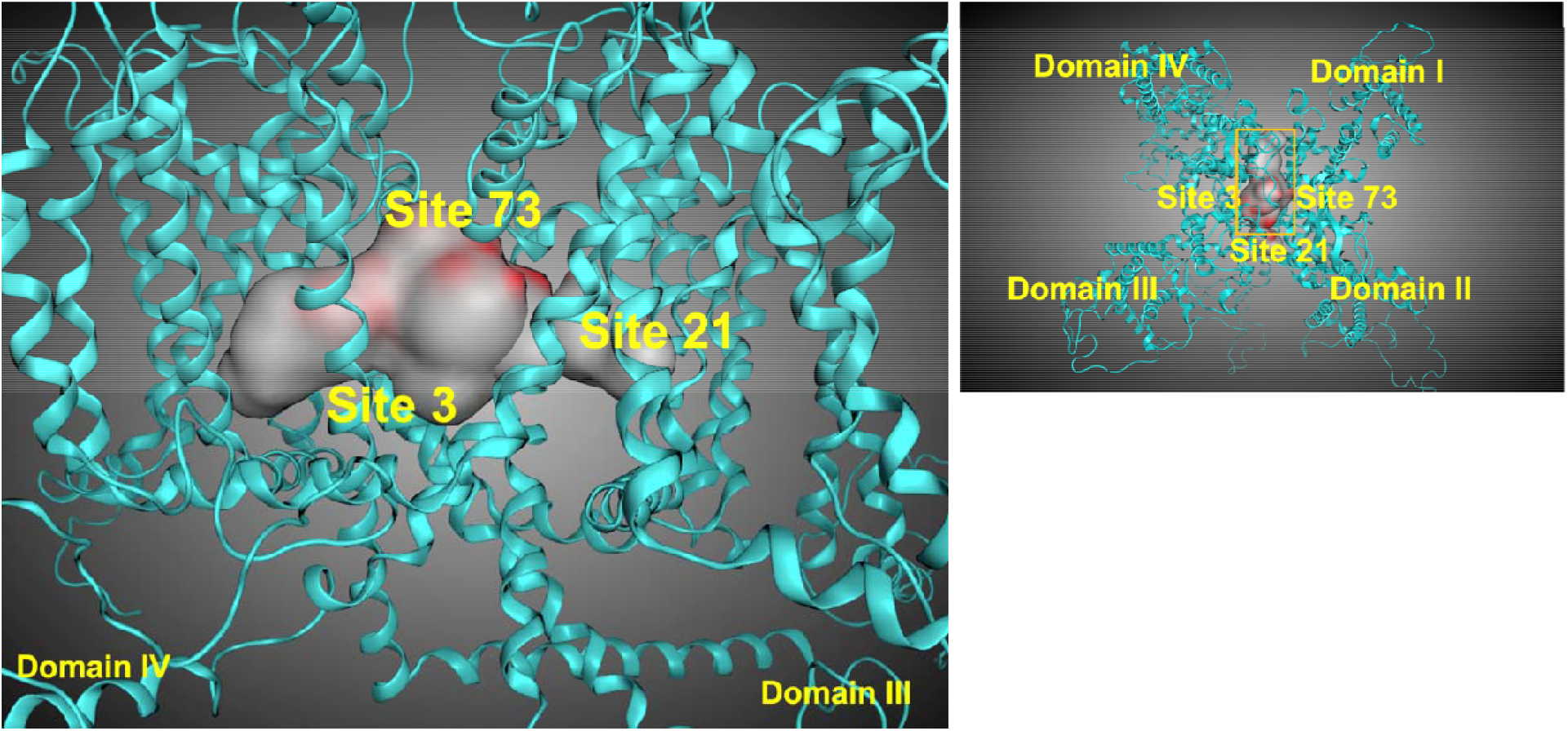
Ligand binding sites in the structure of the constructed hCaV1.2 channel. Three sites were found within the pore of the hCaV1.2 channel. Site numbers were indicated by the order found in the site finder tool.

**SI Table 1:**
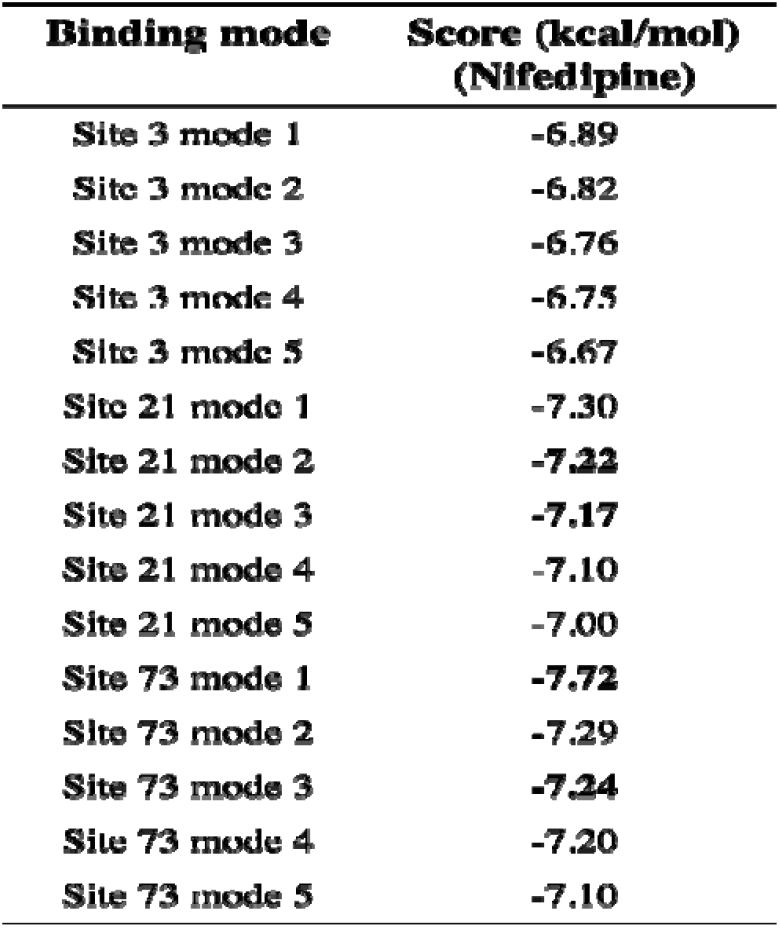
Binding modes with the relative scorings of NIFE.

**SI Figure 3.**
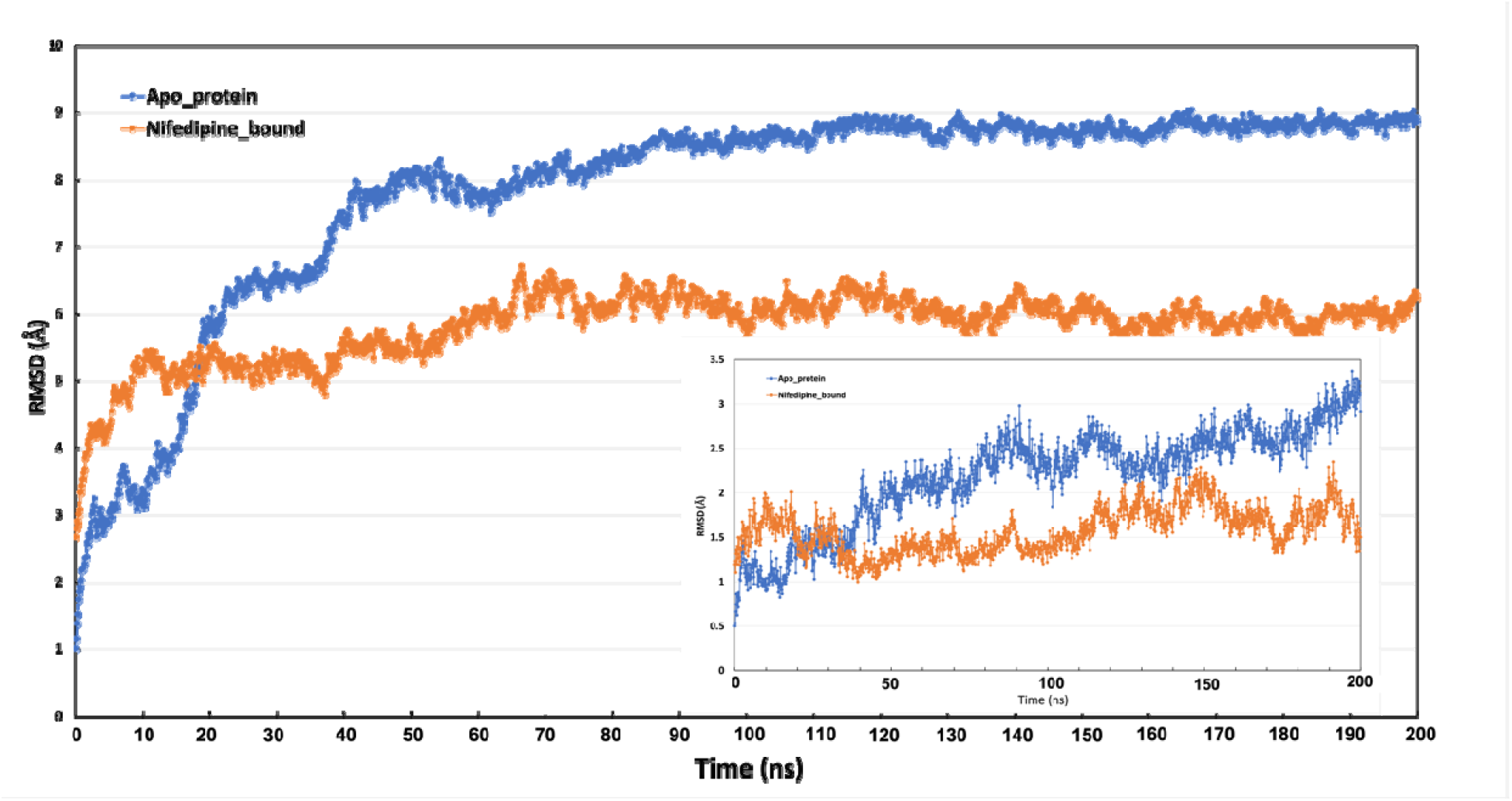
Comparison of the RMSD over 200 ns MD simulations between the Apoprotein and NIFE bound structures. Inlet is the RMSDs of ligand bound pocket.

**SI Figure 4.**
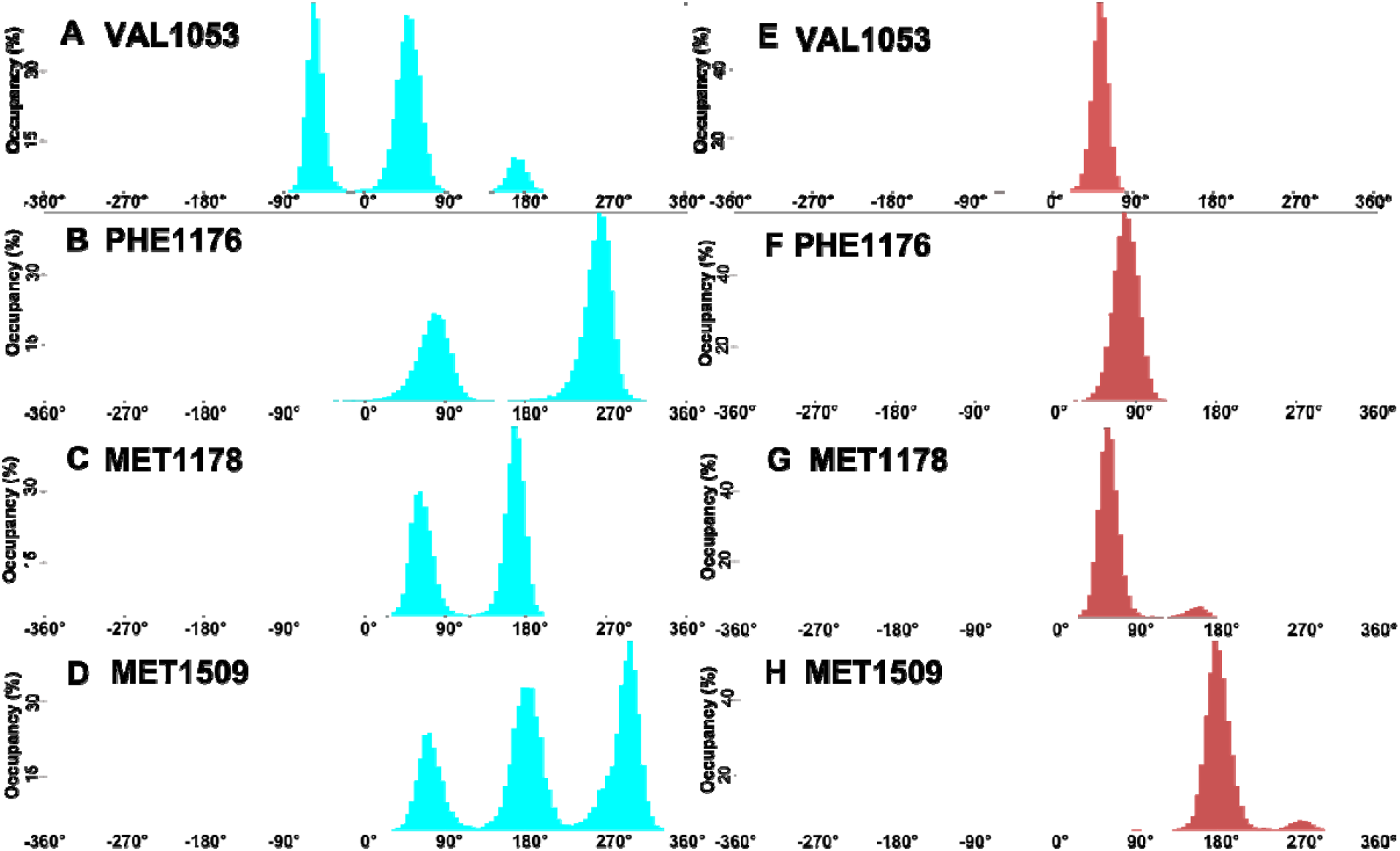
Clustering conformations of residues surrounding the ligand binding pocket. Each histogram can represent one rotameric state. (A)/(E) (Val1053), (B)/(F) (Phe1176), C/G (Met1178), D/H (Met1509). Left panel is for apoprotein structure and right panel is for NIFE bound structure.

